# Protection of Rhesus Macaque from SARS-Coronavirus challenge by recombinant adenovirus vaccine

**DOI:** 10.1101/2020.02.17.951939

**Authors:** Yiyou Chen, Qiang Wei, Ruobing Li, Hong Gao, Hua Zhu, Wei Deng, Linlin Bao, Wei Tong, Zhe Cong, Hong Jiang, Chuan Qin

**Affiliations:** NewHorizon Health Co. Ltd, 13F Building T1, No. 400 Jiang’er Road, Hangzhou Binjiang District, Zhejiang 310051, China; Key Laboratory of Human Disease Comparative Medicine, Chinese Ministry of Health, Beijing Key Laboratory for Animal Models of Emerging and Remerging Infectious Diseases, Institute of Laboratory Animal Science, Chinese Academy of Medical Sciences, 5 Panjiayuan Nanli, Chaoyang District, Beijing 100021, P.R. China; Novartis Institute of Biomedical Research, No. 4218 Jinke Road, Zhangjiang Hi-tech Park, Shanghai 201203, China

**Author notes:** Corresponding authors &.

## Abstract

A recombinant adenovirus vaccine against the SARS Coronavirus (SARS-CoV) was constructed, which contains fragments from the S, N, and Orf8 genes. Rhesus Macaques immunized with the recombinant adenovirus generated antigen-specific humoral and cellular response. Furthermore, the vaccine provided significant protection against subsequent live SARS-CoV challenge. In contrast, three out of four monkeys immunized with placebo suffered severe alveolar damage and pulmonary destruction.

---

The SARS-CoV is a novel coronavirus first identified in 2003^1,2^, which causes severe respiratory distress in patients and can be lethal, especially in people over 65 years old. There are currently no effective drug or treatment for this disease, and the development of a safe and effective vaccine is a high priority for public health. Although several vaccine candidates have already been tested in mice^3,4^ and monkey^5^, clear evidence of protection in a relevant non-human primate model is still lacking.

A bi-cistronic expression cassette was designed to express multiple antigens according to the genome sequence of the BJ01 strain. The gene cassette contains an S1-orf8 fusion and the N gene, separated by an IRES sequence (**Supplemental Figure 1a**, Genbank acession No. AY278488). Recombinant adenovirus containing the expression cassette, SV8000, can express the mRNA transcripts of both genes in vitro (data not shown). And the S1-orf8 fusion protein was predominantly expressed in the culture supernatant as expected (**Supplemental Figure 1b**).

Rhesus Macaques were immunized twice with SV8000 intramuscularly at week 0 and week 4. Antibody response was relatively low in the first 4 weeks, however it rose quickly after the booster injection (**Figure 1a**). By week 8, all immunized monkeys produced high titers of IgG antibody whereas animals in the placebo group only had baseline level. Neutralizing antibody followed similar trend with the high-dose and low-dose group reaching titers of 95+/-65and 83+/-38, respectively (**Figure 1b**).

**Figure 1.**
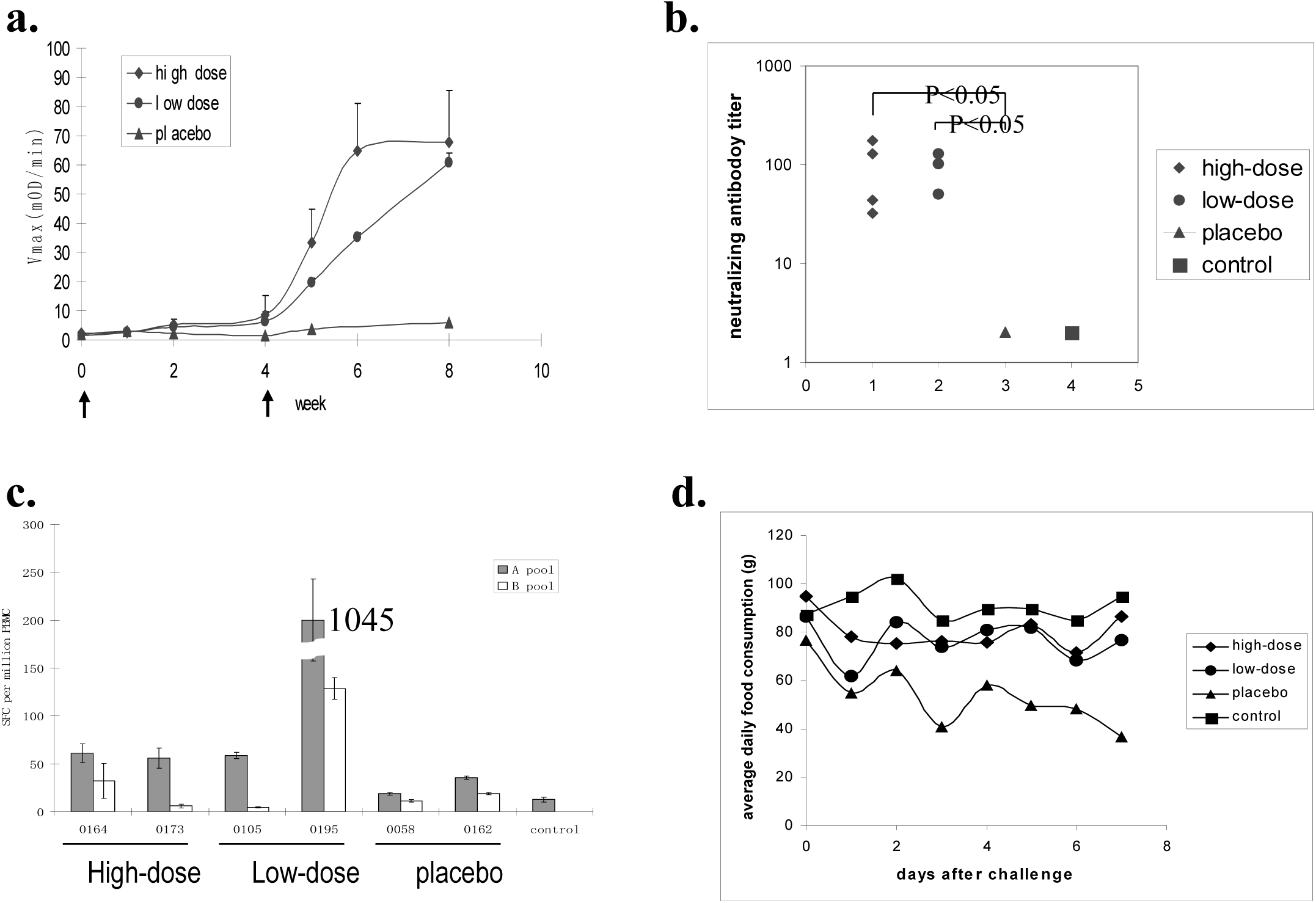
Immune response and physiology of monkeys immunized with recombinant adenovirus vaccine. **a.** Time course of antibody response. Serum samples were diluted 40-fold prior to ELISA analysis. Animals in the high-dose and low-dose group were immunized with 1×10^9^ and 1×10^8^ pfu virus each time, respectively. Arrows indicate the time point for immunization. **b.** Determination of virus neutralization antibody titer using 96-well microneutralization assay. An arbitrate titer of 2 was assigned to animals in the placebo and control group since there was no detectable neutralization activity in these samples. Statistical significance between different data sets was calculated using t-test. **c.** ELISPOT analysis of IFN-γ response in PBMC samples. Pool A and B contain peptides corresponding to S1-orf8 fusion and N, respectively. Data represent two animals from each group. **d.** Average food consumption after live virus challenge.

In order to test whether the recombinant adenovirus can illicit antigen-specific T cell response, pools of overlapping peptides corresponding to target antigen sequences were used to stimulate PBMCs harvested from monkeys four weeks after the booster injection. Animals from immunized groups generated detectable IFN-γ response (**Figure 1c**), whereas the placebo group did not show any significant response above background. Furthermore, peptides from both pools were able to induce IFN-γ release from immunized samples, indicating that both genes were expressed in vivo and were recognized by the immune system. Interestingly, the highest T cell response came from monkey #0195 in the low-dose group, which had at least10-times higher response than any other animal.

To test the protective efficacy of the recombinant adenovirus vaccine, monkeys in the high-dose, low-dose, and placebo group were challenged intranasally with 10^5^TCID_50_ live virus of the PUMC-1 strain (Genbank Accession No. AY350750). Two additional uninfected monkeys housed in the same facility were used as negative control.

Three days after virus inoculation, monkeys in the placebo group became lethargic, and had markedly reduced food intake (**Figure 1d**). Two animals displayed respiratory distress, including shortness of breath and increased respiratory rate. In contrast, none of the immunized animals, or the control animals, displayed any visible respiratory stress during the entire period of the experiment.

Pharyngeal swab and serum samples were collected on day 2, 5, and 7 after challenge. The presence of SARS-CoV was detected via either RT-PCR or viral culture (**Table 1**). Three out of eight immunized animals had detectable RNA, and only one of which was positive on day 7. In contrast, viral RNA could be detected in all four animals in the placebo group, and it persisted for at least 5 days. The absolute viral load also appeared to be significantly higher than that of the immunized animals.

**Table 1.**
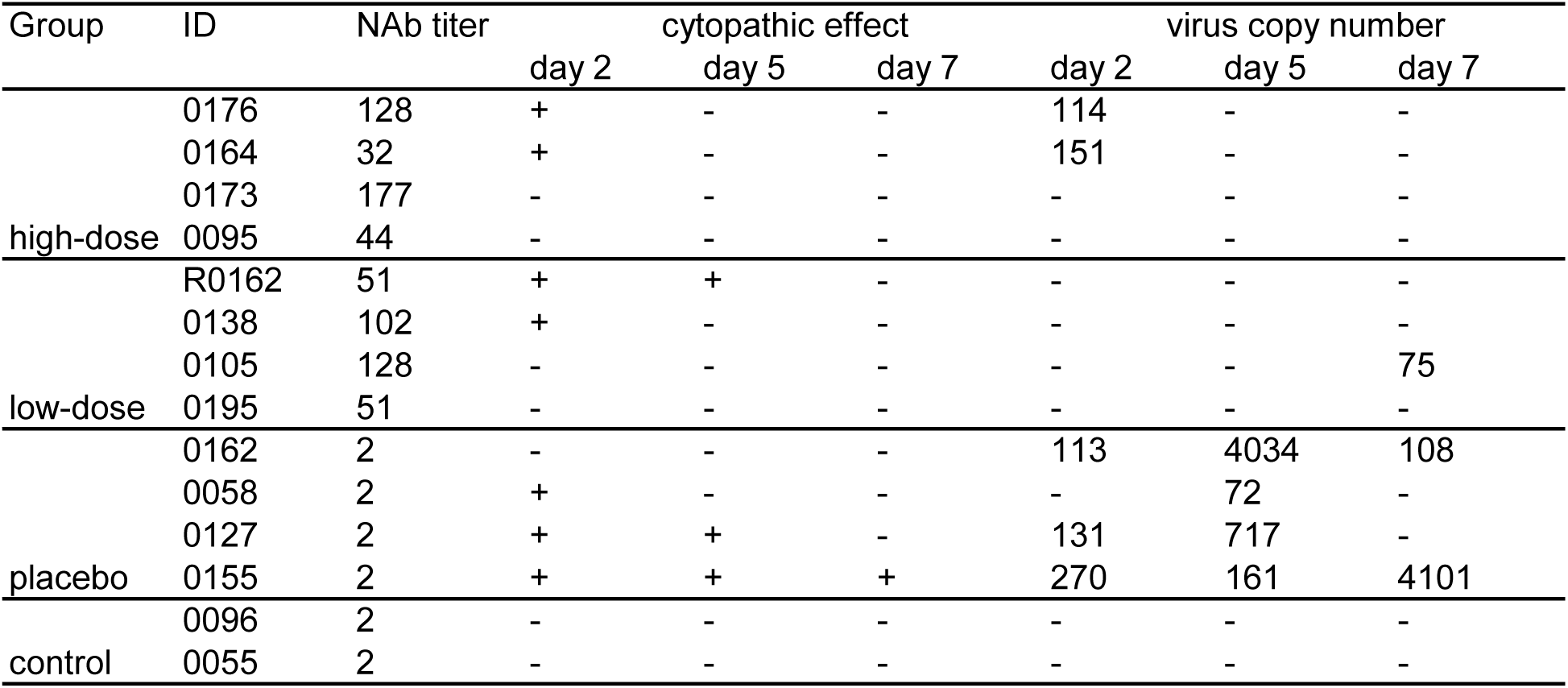
Virus detection of postmortem tissues from Rhesus Macaques infected with the PUMC-1 strain SARS-CoV. Negative value represents no visible cytopathic effect (CPE) after three consecutive passages. Quantitative RT-PCR reaction was used to quantify the presence of viral RNA in the pharyngeal swab samples. The genome copy number was determined according to a standard curve according to the Manufacturer’s manual.

Pharyngeal swab samples were also used to inoculate Vero cells in order to re-isolate live virus after the challenge. Infectious viral particles can be isolated from four out of eight animals in the immunized animals, and only one of which had active viral replication by day 5. In contrast, infectious viral particles can be isolated from three out of four animals in the placebo group, two of which had active viral replication by day 5. Postmortem tissues, including serum, lung, kidney, spleen, liver, and lymph node, were also analyzed, and no positive culture could be identified in any sample. As expected, no infectious viral particles can be isolated from the control animals.

Consistent with previous observations^6^, animals in the placebo group suffered severe pulmonary damage, characterized by extensive disruption of alveoli walls and the accumulation of proteinaceous edema fluid in the alveoli space (**Figure 2c**). The alveoli wall was thickened by heavy infiltration of mononuclear cells, and hyaline membrane could be identified lining the alveoli walls (**Supplemental Figure 2b/c**). In the most severe cases, extensive destruction of the epithelial cell layer and local hemorrhage could also be observed. Histology for the high-dose animals (**Figure 2b**) were overall comparable to that of the control animals (**Figure 2a**), although in some instances minor edma and mononuclear cell infiltration in the alveoli wall could be observed. The phenotype for the low-dose group was somewhat in between the high-dose and the placebo group (**Supplemental Figure 2a**), with pronounced thickening of the alveoli wall in some cases and much less severe pathology in other cases.

**Figure 2.**
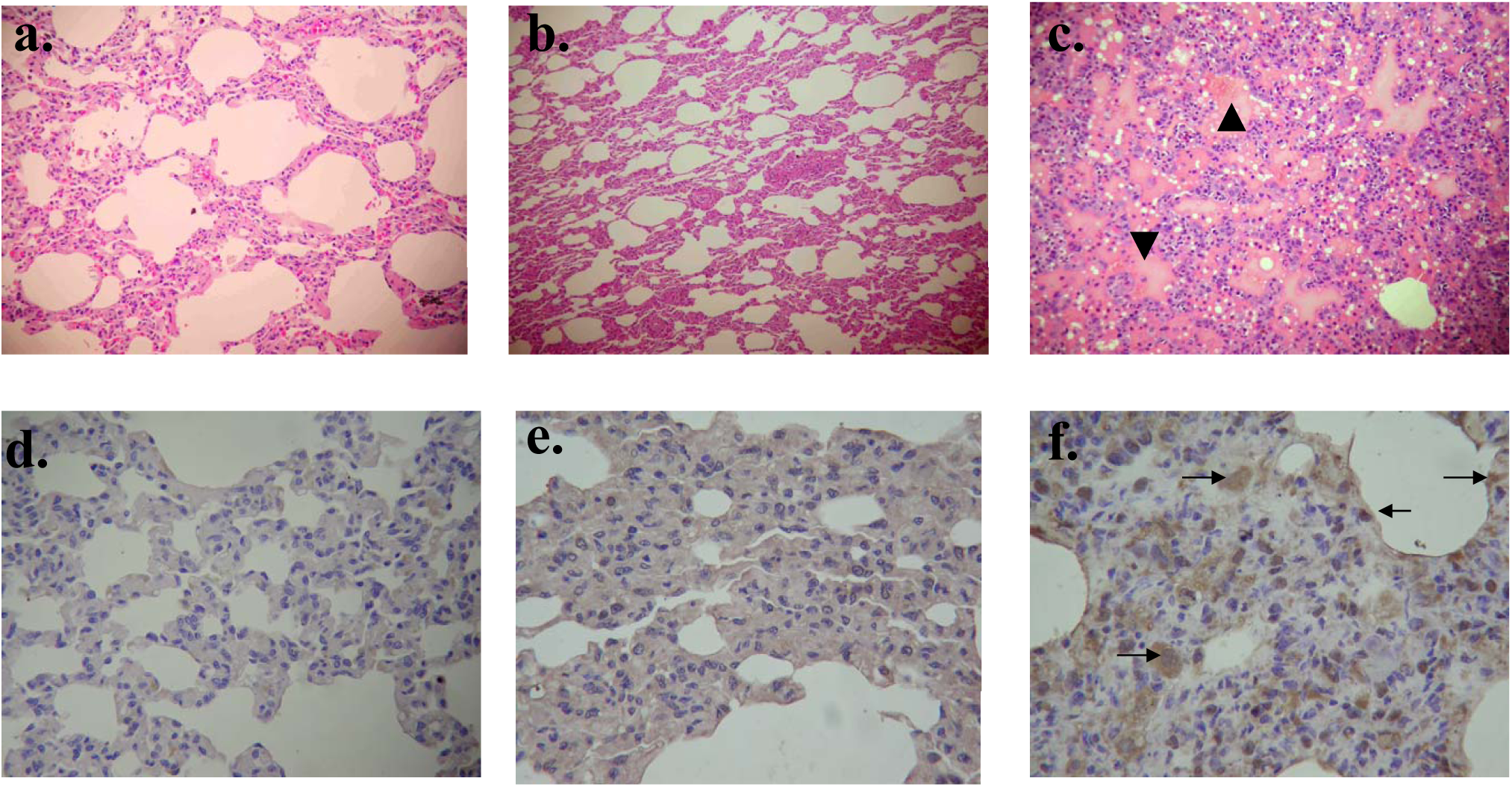
Histology and IHC study on postmortem lung sections from rhesus macaques infected with live SARS-CoV. All animals were euthanized 7 days after viral challenge, and anti-SARS-CoV monoclonal antibody was used to identify SARS antigens in the tissue. Representative data from lung sections were shown. Histology samples include: **a.** Control group. **b.** High-dose group. **c.** Placebo group. The arrowheads point to alveoli space filled with proteinaceous fluid. IHC samples include: **d.** Control group. **e.** High-dose group. **f.** Placebo group. Arrows point to cells that were positive for SARS antigen expression.

In order to correlate the disease pathology with viral replication *in situ*, sections of the lung were also analyzed by immunohistochemistry. There was minimal, if any, staining in the sections for the high-dose group (**Figure 2e**), which was similar to that of control (**Figure 2d**). In contrast, sections for the placebo group showed numerous positive cells, with predominantly cytosolic staining, indicating continuing antigen expression in these cells (**Figure 2f**). The low-dose group also had significant antigen present within the alveoli wall and the epithelial cells (**Supplemental Figure 2d**).

Despite of recent advances, the development of an effective vaccine for SARS-CoV is likely to encounter significant hurdles in the future. One potential roadblock is the emergence of viral mutations^7^. The PUMC-1 strain differs from the BJ01 strain in two positions within the S1 domain (G->D on position 21,721, and I->T on position 22,222). It was speculated that these two sites might be involved in a viral escape mechanism, since all the hotel M associated cases carry D and T whereas other cases in Northern China had G and I, respectively^8,9^. Inclusion of multiple target genes in the vaccine design can potentially reduce the likelihood of viral escape.

Here we have shown that Rhesus Macaques immunized with SV8000, especially those in the high-dose group, had lower viral load, shorter viral persistence period, less severe pathological damages, and importantly almost no visible respiratory stress symptoms. All the data collectively demonstrated the validity of using a recombinant adenovirus vaccine to prevent SARS-CoV infection, and further clinical evaluation in human is warranted.

## Acknowledgements

We are grateful for Dr. Michael Buchmeirer and Professor Binggen Ru for providing critical discussions, and we thank Dr. Xinmin Tu, Dr. Qingyu Zhu, and Dr. Ede Qin for technical assistance. Additional financial support came from the government of Spain and the Mitsui & Co.

## Supplemental figure legends

**Supplemental figure 1.**
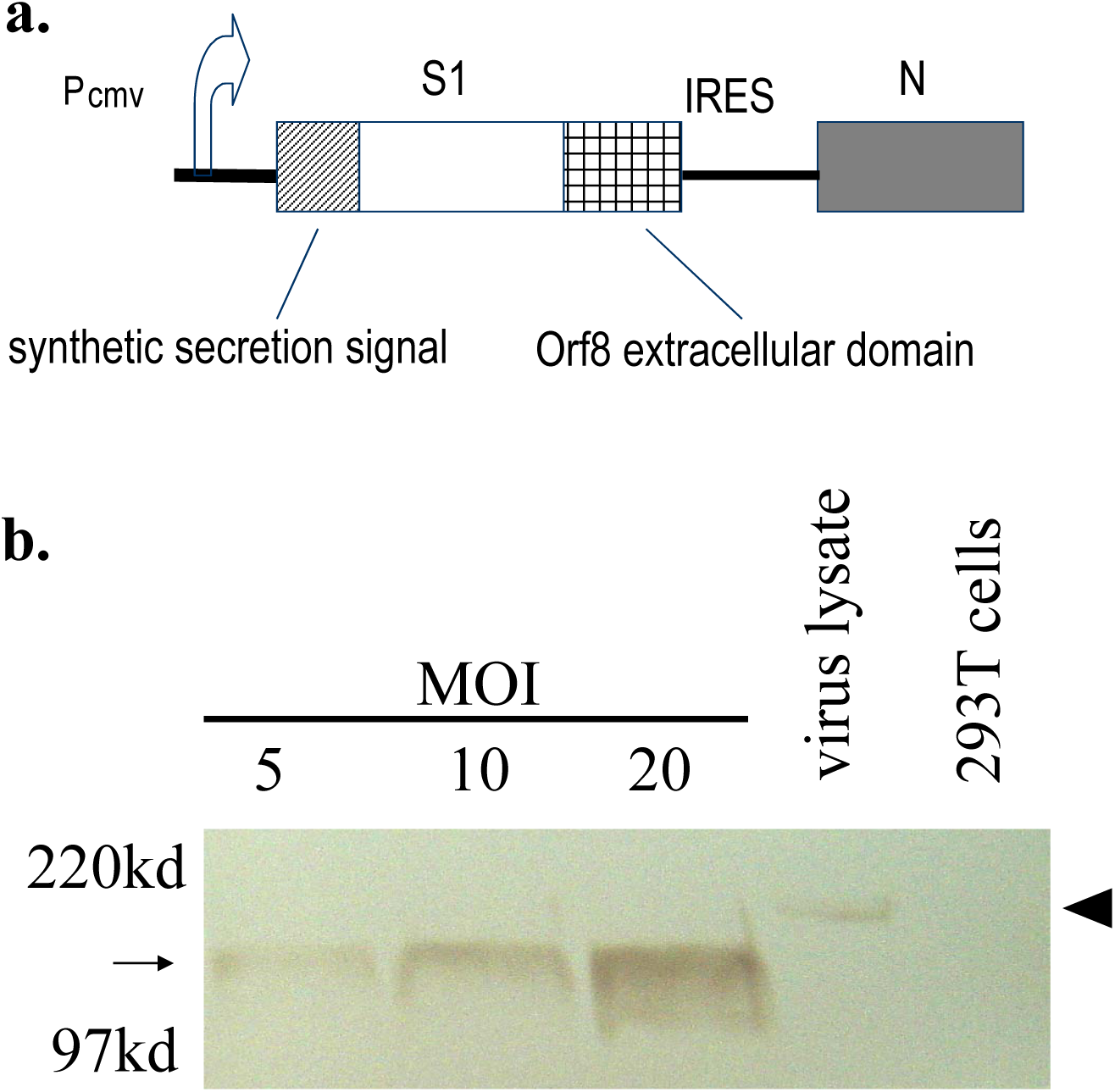
Schematic gene design and protein expression in vitro. **a.** structure of the expression cassette. **b.** Target gene expression in vitro. 293T cells were transduced with recombinant adenovirus, SV8000. Supernatant was harvested 48hr after transduction. S1-orf8 protein was identified using monoclonal antibody against SARS-CoV. SARS-CoV infected Vero E6 cell lysate was used as positive control.

**Supplemental figure 2.**
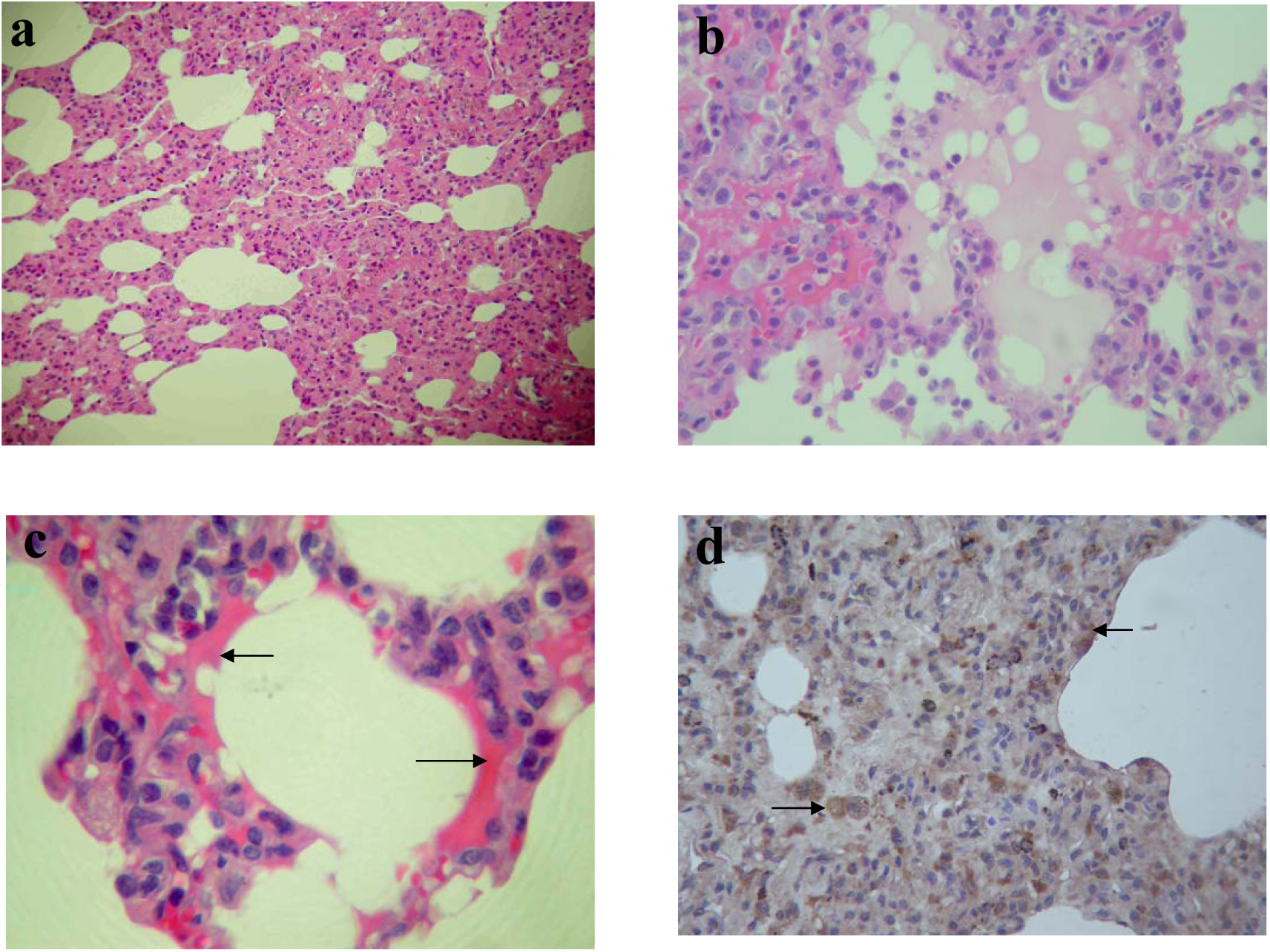
Histology and IHC studies on post-mortem lung tissue. **a.** lung sections from animals immunized with low-dose vaccine. **b/c.** lung sections from animals immunized with high-dose vaccine. **d.** IHC study on sections from animals in the low-dose group. Arrows point to cells positive for SARS antigen expression.

